# Knockdown of GAS5 restores ox-LDL-induced impaired autophagy flux via upregulating miR-26a in human endothelial cells

**DOI:** 10.1101/293324

**Authors:** Weijie Liang, Taibing Fan, Lin Liu, Lianzhong Zhang

## Abstract

**Background:** Oxidized low-density lipoprotein (ox-LDL)-induced endothelial cell (EC) injury and autophagy dysfunction play a vital role in the development of atherosclerosis. LncRNAs have been identified to participate in the regulation of pathogenesis of atherosclerosis. However, it remains largely undefined whether growth-arrest specific transcript 5 (GAS5) could influence ox-LDL-induced autophagy dysfunction in ECs.

**Methods:** The expressions of GAS5 and miR-26a in the plasma samples of patients with atherosclerosis and ox-LDL-treated human aortic endothelial cells (HAECs) were detected by qRT-PCR. Luciferase reporter assay, RNA immunoprecipitation (RIP), and RNA pull down were performed to validate whether GAS5 could directly interact with miR-26a. The effects of ox-LDL, GAS5 or combined with miR-26a on apoptosis and autophagy were evaluated by flow cytometry analysis and western blot, respectively.

**Results:** GAS5 expression was upregulated and miR-26a was downregulated in the plasma samples of patients with atherosclerosis and ox-LDL-treated HAECs. There was reciprocal inhibition between GAS5 and miR-26a expressions in ox-LDL-treated HAECs. We further demonstrated that GAS5 directly interacted with miR-26a in ox-LDL-treated HAECs. Additionally, ox-LDL administration induced apoptosis and impaired autophagy flux in HAECs. Rescue experiments demonstrated that GAS5 knockdown restored ox-LDL-induced impaired autophagy flux by upregulating miR-26a in HAECs.

**Conclusion:** Knockdown of GAS5 restores ox-LDL-induced impaired autophagy flux via upregulating miR-26a in human endothelial cells, revealing a novel regulatory mechanism for ox-LDL-induced impaired autophagy flux in ECs through ceRNA crosstalk.

## 1 Introduction

Atherosclerosis, a devastating and chronic multi-factorial vascular cardiovascular disease, remains a leading health issue among aged people, accounting for the high morbidity and morbidity of cardiovascular disease worldwide [1]. It has been suggested that endothelial dysfunction is the requisite for the progression of atherosclerosis [2]. Oxidized low-density lipoprotein (ox-LDL) is considered as a key risk factor associated with endothelial dysfunction in atherosclerosis [3]. Extensive researches within the past decades have demonstrated that ox-LDL-induced autophagy dysfunction in endothelial cells (ECs) plays a vital role in the development of atherosclerosis [4]. Autophagy is well-known as a dynamic process of recycling that plays a prominent role in degrading dysfunctional or damaged proteins or intracellular organelles in eukaryotic cells [5]. Recently, increasing evidence has indicated that impaired autophagy flux contributes to lipid metabolism dysfunction and EC apoptosis, greatly implicated in vascular endothelial dysfunction and atherosclerotic plaque development [6]. Therefore, upregulation of autophagy may be a promising therapy to protect ECs from ox-LDL-induced injury.

Recently, increasing evidence demonstrated that noncoding RNAs (ncRNAs), including the recently acknowledged long noncoding RNA (lncRNA) and the well-known microRNA (miRNA), play important roles in the regulation of gene expression via multiple mechanisms [7]. LncRNAs are generally defined as a group of RNA transcripts longer than 200 nucleotides with limited or no protein-coding potential. A wide range of documents unveil that lncRNAs play important functional roles in the regulation of lipid metabolism, vascular inflammation, cell proliferation, and EC apoptosis, suggesting that lncRNAs participate in the regulation of pathogenesis of atherosclerosis [8]. LncRNA growth-arrest specific transcript 5 (GAS5), located at chromosome 1q25.1, was originally isolated from mouse NIH 3T3 cells using subtraction hybridization [9]. There is striking evidence that GAS5 functions as a tumor suppressive lncRNA and is aberrantly downregulated in a variety of human cancers [10]. Notably, previous studies showed that GAS5 was increased in the plaque of atherosclerosis collected from patients and animal models and knockdown of GAS5 reduced the apoptosis of macrophages and vascular endothelial cells after ox-LDL stimulation [11, 12]. However, it remains largely undefined whether GAS5 could influence ox-LDL-induced autophagy dysfunction in ECs.

miRNAs are small, endogenous, single-stranded ncRNAs with 20-25 nucleotides in length, which repress gene expression at the posttranscriptional level via mRNA degradation or translational inhibition. More recently, substantive studies have demonstrated that miRNAs play critical roles in the development of atherosclerosis via regulating the proliferation, migration and apoptosis of various types of cells [13]. miR-26a, a highly conserved miRNA, has been revealed to play essential roles in development, cell differentiation, apoptosis and growth. miR-26a is reported to be frequently dysregulated in cardiovascular diseases such as cardiac hypertrophy and myocardial ischemia [14, 15]. Moreover, it was previously reported that miR-26a was downregulated in high-fat diet (HFD)-fed apolipoprotein E (apoE)^−/-^ mice and ox-LDL-stimulated human aortic endothelial cells (HAECs) and miR-26a overexpression alleviated the development of atherosclerosis [16, 17].

Recently, a new regulatory mechanism has been proposed that lncRNAs function on a competitive endogenous RNA (ceRNA) to regulate the expressions and biological function of miRNAs [18]. The lncRNA-miRNA interaction has been identified in various human diseases, including vascular pathophysiology [19]. Since our bioinformatics analysis demonstrated that GAS5 contained the complementary binding sites in miR-26a, we hypothesized whether GAS5 could function as a molecular sponge of miR-26a to regulate atherosclerosis progression.

In the present study, we investigated the effects of GAS5 on ox-LDL-induced autophagy in HAECs, as well as the interaction between GAS5 and miR-26a.

## 2 Materials and methods

### 2.1 Clinical specimens

A total of 30 atherosclerotic patients diagnosed by clinical symptoms and coronary angiography at the department of cardiology in the Henan Provincial People’s Hospital between January 2015 and August 2016 and 30 healthy subjects were enrolled in the present study. The plasma samples were collected from all patients and healthy subjects and stored at −80°C for further experiments. The study was performed with the approval of the Research Ethics Committee of Henan Provincial People’s Hospital and written informed consent was obtained from each participant.

### 2.2 Cell culture and transfection

HAECs were obtained from American Type Culture Collection (ATCC, Manassas, VA, USA) and were cultured in endothelial cell medium containing endothelial cell growth factors, 10% heat-inactivated fetal bovine serum (FBS, Hyclone, Logan, UT, USA) and 100 U/ml penicillin and 100 µg/ml streptomycin (Sangon, Shanghai, China) at 37°C in a humidified atmosphere of 5% CO_2_. Cells in logarithmic phase were collected for further experiments. When grown to 70-80% confluence, HAECs were transiently transfected with miR-26a mimic (miR-26a), miRNA negative control (miR-NC), miR-26a inhibitor (anti-miR-26a), inhibitor negative control (anti-miR-NC), pcDNA-GAS5 (GAS5), pcDNA empty control (pcDNA), siRNA against GAS5 (si-GAS5), siRNA negative control (si-NC) (all from RiboBio Co., Ltd., Guangzhou, China) using Lipofectamine 2000 reagent (Invitrogen, CA, Carlsbad, USA). Following 48 h of transfection, HAECs were treated with different concentrations of ox-LDL (0.25, 0.5, and 1 mg/L) (UnionBiol, Beijing, China) for 24 h.

### 2.3 Quantitative real-time PCR

Total RNA was extracted from clinical samples or treated HAECs using TRIzol reagent (Invitrogen) and complementary DNA (cDNA) was synthesized from 1 μg of total RNA using a High-Capacity cDNA Reverse Transcription kit (Applied Biosystems, Foster City, CA, USA). The expressions of GAS5 and miR-26a were examined using a SYBR Premix Ex Taq II (Takara, Dalian, China) and TaqMan MicroRNA Assay Kit (Applied Biosystems) on a CFX96 real-time PCR System (Bio-Rad, Hercules, CA, USA), respectively. The expressions of GAS5 and miR-26a were normalized to GAPDH and U6 small nuclear RNA (snRNA), respectively. The relative gene expression was calculated using the 2^−ΔΔCt^ method.

### 2.4 Western blot

Total proteins were extracted from the treated HAECs using RIPA buffer containing protease inhibitors (Roche, Nutley, NJ, USA). Equal quantities of protein were fractionated by a 10% sodium dodecyl sulfate-polyacrylamide gel (SDS-PAGE) and transferred to a nitrocellulose membranes (Millipore, Billerica, MA, USA). After being blocked in 5% skimmed milk in Tris-based saline with Tween 20 (TBST) for 1 h at room temperature, the membranes were incubated with primary antibodies against LC3-II (1:1000, Abcam, Cambridge, UK), LC3-I (1:1000, Abcam), p62 (1:2000, Cell Signaling Technology, Danvers, MA, USA). After washing with TBST, the membranes were incubated with HRP-conjugated secondary antibody for 2 h at room temperature. The protein bands were visualized using a chemiluminescence kit (GE Healthcare, Buckinghamshire, UK) and protein intensity was quantified with Image-Pro Plus 6.0 software (GE Healthcare).

### 2.5 Flow cytometry analysis

Cell apoptosis was assessed using a FITC-Annexin V Apoptosis Detection Kit (BD Bioscience, San Jose, CA, USA). HAECs were seeded into 24-well plates and exposed to ox-LDL at a series of concentration (0.25, 0.5, and 1 mg/L) for 24 h, or transfected with si-NC, si-GAS5, si-GAS5 + anti-miR-NC, si-GAS5 + anti-miR-26a, followed by stimulation with 1 mg/L ox-LDL. Then cells were collected and digested using 0.25% trypsin. After washed twice with ice-cold PBS, cells were resuspended with 500 μl of 1 × binding buffer solution and incubated with 5 μl of Annexin-V FITC and 5 μl propidium iodide (PI) for 15 min in the dark. Apoptotic cells were measured by a BD FACSCalibur flow cytometer (Beckman Coulter, Fullerton, CA, USA) and analyzed with FlowJo software (TreeStar, Ashland, OR, USA).

### 2.6 Luciferase reporter assay

The wild-type fragment of GAS5 including the miR-26a binding sites and its mutant sequence were subcloned into psiCHECK-2 luciferase reporter vector (Promega, Madison, WI, USA) and named as GAS5-WT and GAS-MUT. For luciferase reporter assay, HAECs were cotransfected with 100 ng constructed luciferase reporter vectors, together with 20 ng Renilla luciferase vector, and 100 nM miR-26a or miR-NC using Lipofectamine 2000 (Invitrogen), followed by treatment with 1 mg/L ox-LDL. At a point of 48 h transfection, cells were harvested and the luciferase activity was detected using the dual-Luciferase Reporter Assay System (Promega). The relative luciferase activity was normalized to the Renilla luciferase activity.

### 2.7 RNA immunoprecipitation (RIP) assay

Magna RIP^TM^ RNA-Binding Protein Immunoprecipitation Kit (Millipore, Bedford, MA, USA) was used to perform RIP assay was conducted using the. Briefly, ox-LDL-treated HAECs at 80-90% confluence were collected and lysed in complete RIP lysis buffer. Next, the supernatant from cell lysates was harvested by centrifugation and then 100 μl of cell extracts were incubated with RIP buffer containing A + G magnetic beads conjugated with human anti-Argonaute2 (Ago2) antibody (Abcam) or corresponding negative control IgG (Abcam). To remove the non-specific binding, the Sepharose beads were incubated with Proteinase K to digest the protein and subsequently the precipitated RNA was isolated by TRIzol reagent (Invitrogen). The purified RNA was subjected to qRT-PCR analysis.

### 2.8 RNA pull-down assay with biotinylated GAS5

Briefly, biotinylated DNA probe complementary to GAS5 was amplified by PCR with a T7-containing primer and cloned into the plasmid vector GV394 (Genechem, Shanghai, China). The resultant plasmids were linearized with *Xho*I. Biotin-labeled RNAs were then reversely transcribed with the Biotin RNA Labeling Mix (Roche, Indianapolis, IN, USA) and T7 RNA polymerase (Roche), treated with RNase-free DNase I (Roche) and purified with the RNeasy Mini Kit (Qiagen, Inc., Valencia, CA, USA). The bound RNAs were extracted for further evaluation by qRT-PCR analysis.

### 2.9 RNA pull-down assay with biotinylated miR-26a

Biotinylated miR-26a (bio-miR-26a-WT), biotinylated mutant (bio-miR-26a-MUT), and biotinylated NC (bio-NC) were synthesized by GenePharma (Shanghai, China). HAECs were transfected with biotinylated miRNA using Lipofectamine 2000 (Invitrogen) and collected at 48 post-transfection. Cell lysates were incubated with M-280 streptaviden magnetic beads (Invitrogen). The bound RNAs were purified using TRIzol reagent (Invitrogen) for further qRT-PCR analysis.

### 2.10 Statistical analysis

The data are presented as the mean ± standard deviation (SD). All statistical analyses were performed using SPSS 17.0 software (SPSS, Chicago, IL, USA). The significance of differences was estimated by Student’s *t* test or one-way analysis of variance (ANOVA). Differences were considered to be statistically significant when *P* value < 0.05.

## 3 Results

### 3.1 GAS5 expression was upregulated and miR-26a was downregulated in the plasma samples of patients with atherosclerosis and ox-LDL-treated HAECs

The expressions of GAS5 and miR-26a in the plasma samples from atherosclerotic patients and healthy controls were firstly detected by qRT-PCR. The results indicated that GAS5 expression was abnormally higher while miR-26a expression was aberrantly lower in the plasma samples from patients with atherosclerosis compared with that from healthy controls (Fig. 1A and 1B). Next, we further analyzed the expressions of GAS5 and miR-26a in HAECs treated with different concentrations of ox-LDL (0.25, 0.5, and 1 mg/L). qRT-PCR analysis demonstrated that ox-LDL stimulation dose-dependently increased GAS5 expression and decreased miR-26a expression in HAECs (Fig. 1C and 1D). Collectively, these data suggested that abnormally expressed GAS5 and miR-26a may be associated with the development of atherosclerosis.

**Figure 1.**
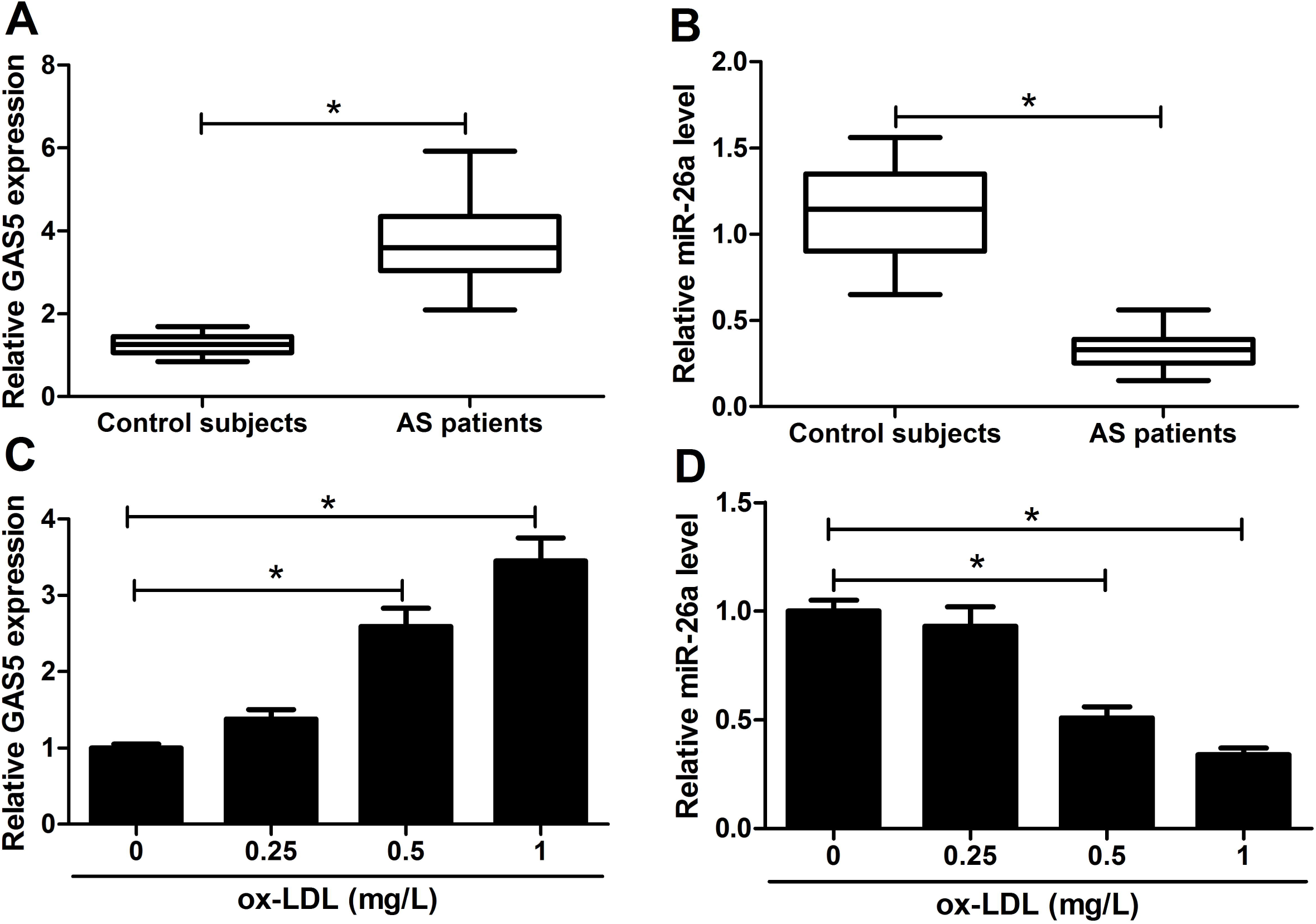
GAS5 expression was upregulated and miR-26a was downregulated in the plasma samples of patients with atherosclerosis and ox-LDL-treated HAECs. qRT-PCR analysis of GAS5 (A) and miR-26a (B) in the plasma samples from patients with atherosclerosis and healthy subjects. qRT-PCR analysis of GAS5 (C) and miR-26a (D) in the HAECs treated with different doses of ox-LDL (0.25, 0.5, and 1 mg/L) for 24 h. \**P* < 0.05.

### 3.2 The relationship between GAS5 and miR-26a expression in ox-LDL-treated HAECs

To address the interaction between GAS5 and miR-26a, HAECs were transfected with GAS5, si-GAS5, miR-26a, anti-miR-26a, or respective controls, followed by ox-LDL stimulation. We found that GAS5 expression was strikingly elevated by GAS5 transfection and greatly reduced by si-GAS5 introduction in ox-LDL-treated HAECs (Fig. 2A). However, ox-LDL-treated HAECs transfected with GAS5 showed a remarkable decline of miR-26a expression and ox-LDL-treated HAECs introduced with si-GAS5 exhibited a substantial enhancement of miR-26a expression (Fig. 2B). Besides, miR-26a expression was considerably enhanced following transfection of miR-26a but dramatically reduced after treatment with anti-miR-26a in ox-LDL-treated HAECs (Fig. 2C). On the contrary, GAS5 expression was obviously lowered in HAECs treated with miR-26a and ox-LDL but evidently augmented in HAECs treated with anti-miR-26a and ox-LDL (Fig. 2D). Together, these findings uncovered that there was reciprocal inhibition between GAS5 and miR-26a expressions in ox-LDL-treated HAECs.

**Figure 2.**
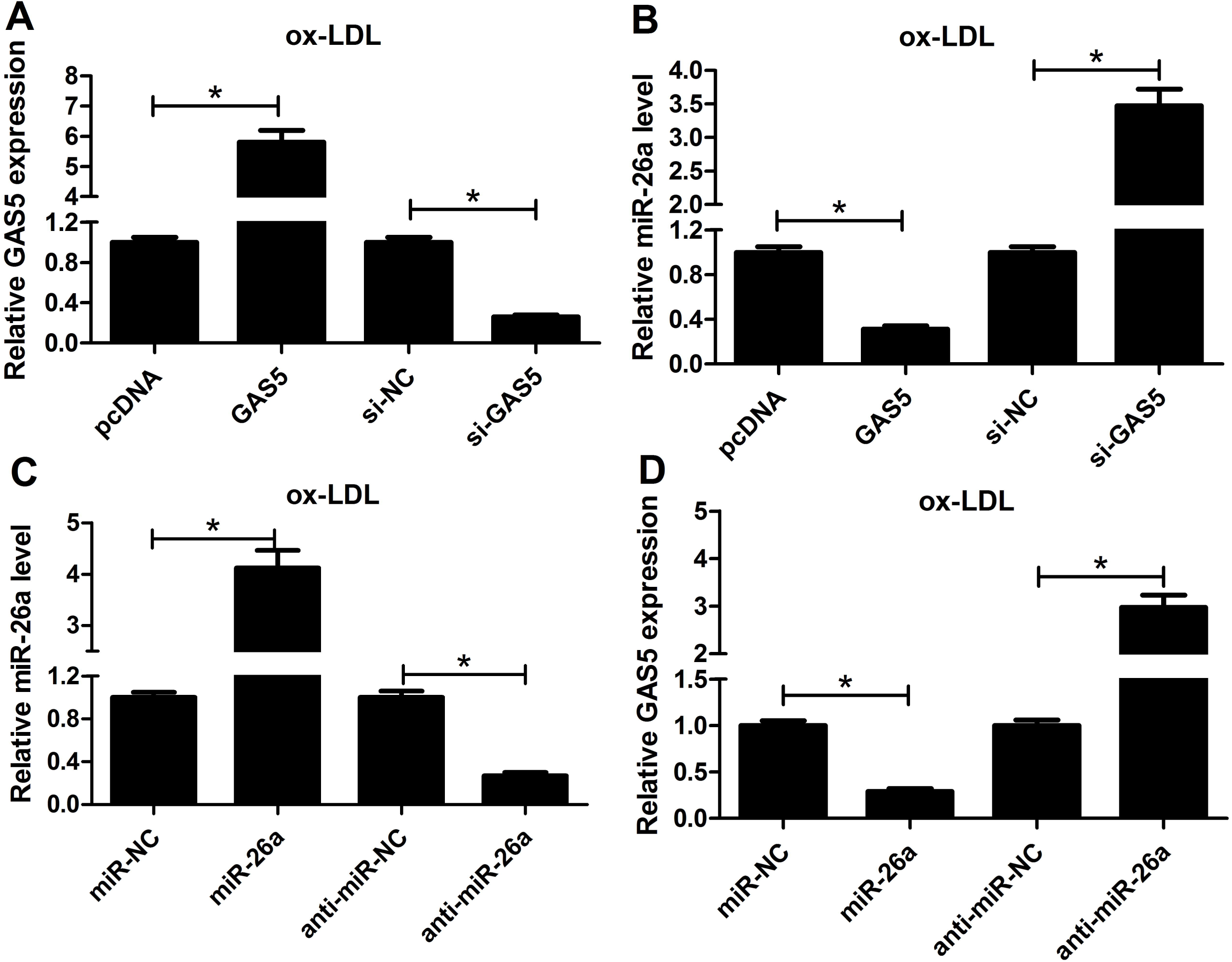
The relationship between GAS5 and miR-26a expression in ox-LDL-treated HAECs. (A and B) The expressions of GAS5 and miR-26a were examined by qRT-PCR in HAECs after transfection with GAS5, si-GAS5, or corresponding controls, followed by administration with 1 mg/L ox-LDL for 24 h. (C and D) The expressions of GAS5 and miR-26a were examined by qRT-PCR in HAECs after introduction with miR-26a, anti-miR-26a, or matched controls, followed by administration with 1 mg/L ox-LDL for 24 h. \**P* < 0.05.

### 3.3 GAS5 directly interacted with miR-26a in ox-LDL-treated HAECs

Based on the reciprocal repression between GAS5 and miR-26a expression, we guessed whether the crosstalk between GAS5 and miR-26a is through direct interaction. Accordingly, bioinformatics analyses were performed to predict the potential miRNAs for GAS5. As a result, the prediction showed the potential binding sites for miR-26a on GAS5, as displayed in Fig. 3A. To verify whether GAS5 could directly interact with miR-26a, we cloned the wild-type or mutated fragment of GAS5 into psiCHECK-2 luciferase reporter vector and performed luciferase reporter assay. The results manifested that cotransfection with GAS5-WT and miR-26a significantly decreased the luciferase activity, but cotransfection with GAS5-MUT and miR-26a did not affect the luciferase activity in ox-LDL-stimulated HAECs (Fig. 3B). It is reported that miRNAs exerted their gene silencing functions via binding to Ago2, a core component of the RNA-induced silencing complex (RISC) [20]. Then RIP assay was performed in cell extracts from HAECs treated with utilizing the antibody against Ago2 and the results presented that GAS5 and miR-26a were both preferentially enriched in Ago2 pellets compared with control IgG immunoprecipitates (Fig. 3C). To confirm whether miR-26a could directly interact with GAS5, we applied a biotin-labeled miRNA pull down to capture GAS5 using M-280 streptaviden magnetic beads from HAECs transfected with biotinylated miR-26a and the results suggested that GAS5 was pulled down as analyzed by qRT-PCR, while miR-26a-MUT with mutant binding sites of GAS5 led to the inability of miR-26a to pull down GAS5 (Fig. 3D). Also, we used the opposite pull down system to confirm whether GAS5 could pull down miR-26a using a biotin-labeled specific GAS5 probe. We observed a significant amount of miR-26a in the GAS5 pulled down pellet compared with control group as analyzed by qRT-PCR (Fig. 3D). These results demonstrated that GAS5 could directly bind to miR-26a in ox-LDL-treated HAECs.

**Figure 3.**
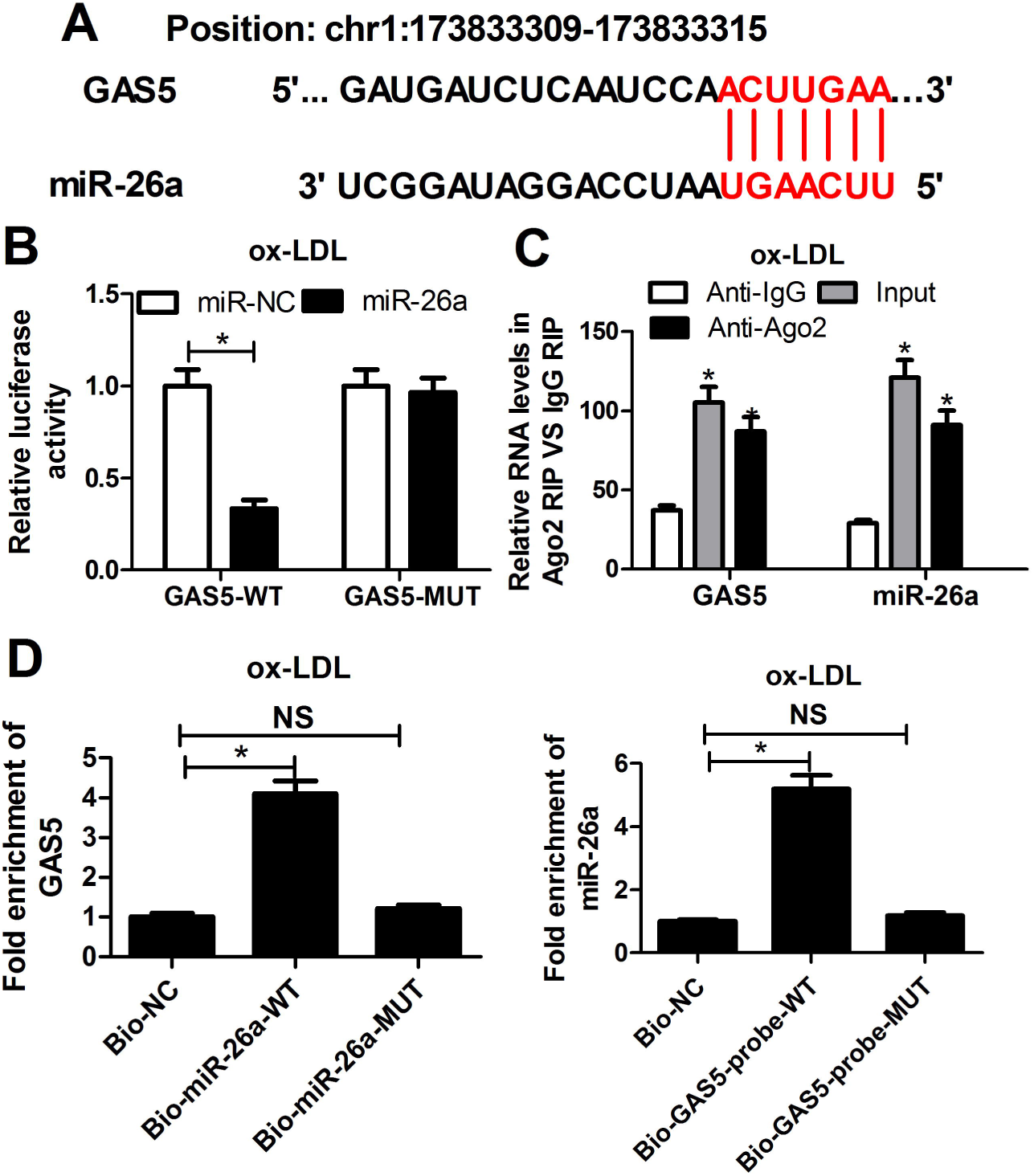
GAS5 directly interacted with miR-26a in ox-LDL-treated HAECs. (A) Predict binding sites between GAS5 and miR-26a. (B) Luciferase reporter assay was performed to detect the luciferase activity in HAECs after cotransfection with GAS5-WT or GAS5-MUT and miR-26a or miR-NC, followed by treatment with ox-LDL. (C) Anti-Ago2 RIP assay was performed in cellular lysates from HAECs treated with ox-LDL to explore the association between GAS5 and miR-26a. HAECs were transfected with biotinylated miR-26a (bio-miR-26a-WT), biotinylated mutant (bio-miR-26a-MUT), and biotinylated NC (bio-NC), followed by treatment with ox-LDL, and collected at 48 post-transfection for pull-down assay with biotinylated miR-26a. Detection of miR-26a using qRT-PCR in the samples pulled down by biotinylated GAS5 and NC probe. \**P* < 0.05.

### 3.4 ox-LDL administration induced apoptosis and impaired autophagy flux in HAECs

Next, we analyzed the effects of ox-LDL on the apoptosis of HAECs. HAECs were treated with different concentrations of ox-LDL (0.25, 0.5, and 1 mg/L) and then flow cytometry analysis was conducted. As shown in Fig. 4A, the percentage of apoptotic rate was specifically increased following ox-LDL treatment in a dose-dependent manner. Moreover, we further evaluated the influence of ox-LDL on the autophagy flux in HAECs by detecting the protein levels of autophagy markers LC3 and p62. The western blot analysis indicated that the LC3-II/LC3-I ratio was distinctly decreased while p62 expression was notably increased in ox-LDL-treated HAECs dose-dependently, suggesting that ox-LDL induced impaired autophagy flux in HAECs (Fig 4B). Taken together, these results demonstrated that ox-LDL administration induced apoptosis and weakened autophagy in HAECs.

**Figure 4.**
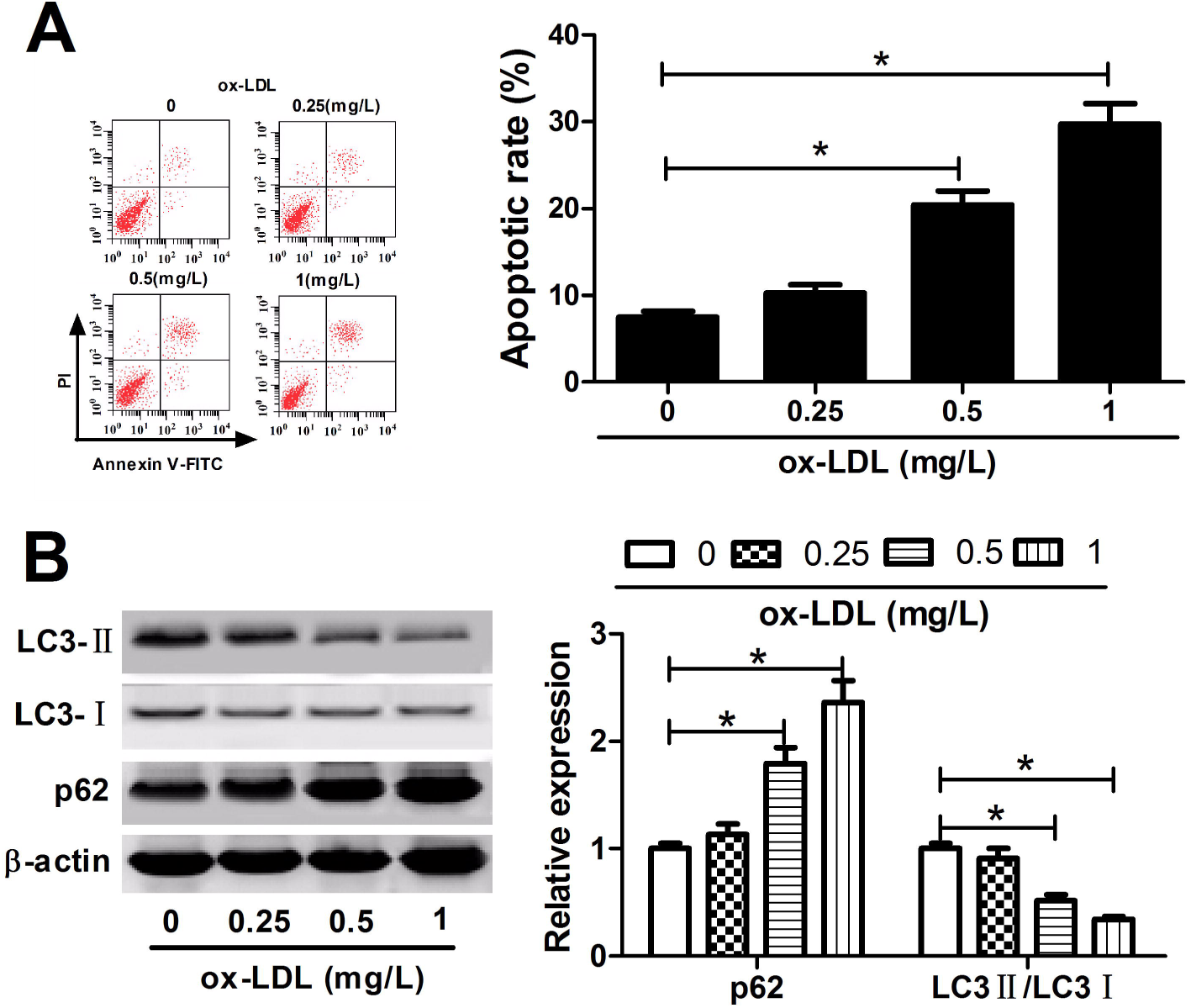
ox-LDL administration induced apoptosis and weakened autophagy flux in HAECs. (A) Flow cytometry analysis was performed to assess the apoptosis of HAECs treated with different concentrations of ox-LDL (0.25, 0.5, and 1 mg/L). (B) Western blot was conducted to determine the protein levels of LC3-II, LC3-I and p62 in HAECs treated with different doses of ox-LDL (0.25, 0.5, and 1 mg/L). *P < 0.05.

### 3.5 GAS5 knockdown restored ox-LDL-induced impaired autophagy flux by upregulating miR-26a in HAECs

To figure out the effects of GAS5 or combined with miR-26a on ox-LDL-induced impaired autophagy flux, HAECs were transfected with si-GAS5, si-NC, si-GAS5 + anti-miR-26a, si-GAS5 + anti-miR-NC, and then exposed to 1 mg/L ox-LDL. Flow cytometry analysis proved that silencing of GAS5 effectively suppressed ox-LDL-induced apoptosis in HAECs, which was partially recuperated by inhibition of miR-26a (Fig. 5A). The subsequent western blot analysis displayed that transfection with si-GAS5 obviously boosted the LC3-II/LC3-I ratio and reduced p62 level in ox-LDL-treated HAECs (Fig. 5B). However, these effects were significantly reversed by anti-miR-26a treatment (Fig. 5B), suggesting that miR-26a suppression attenuated GAS5 knockdown-mediated restoration of ox-LDL-induced impaired autophagy flux in HAECs. These results manifested that GAS5 knockdown restored ox-LDL-induced impaired autophagy flux by upregulating miR-26a in HAECs.

**Figure 5.**
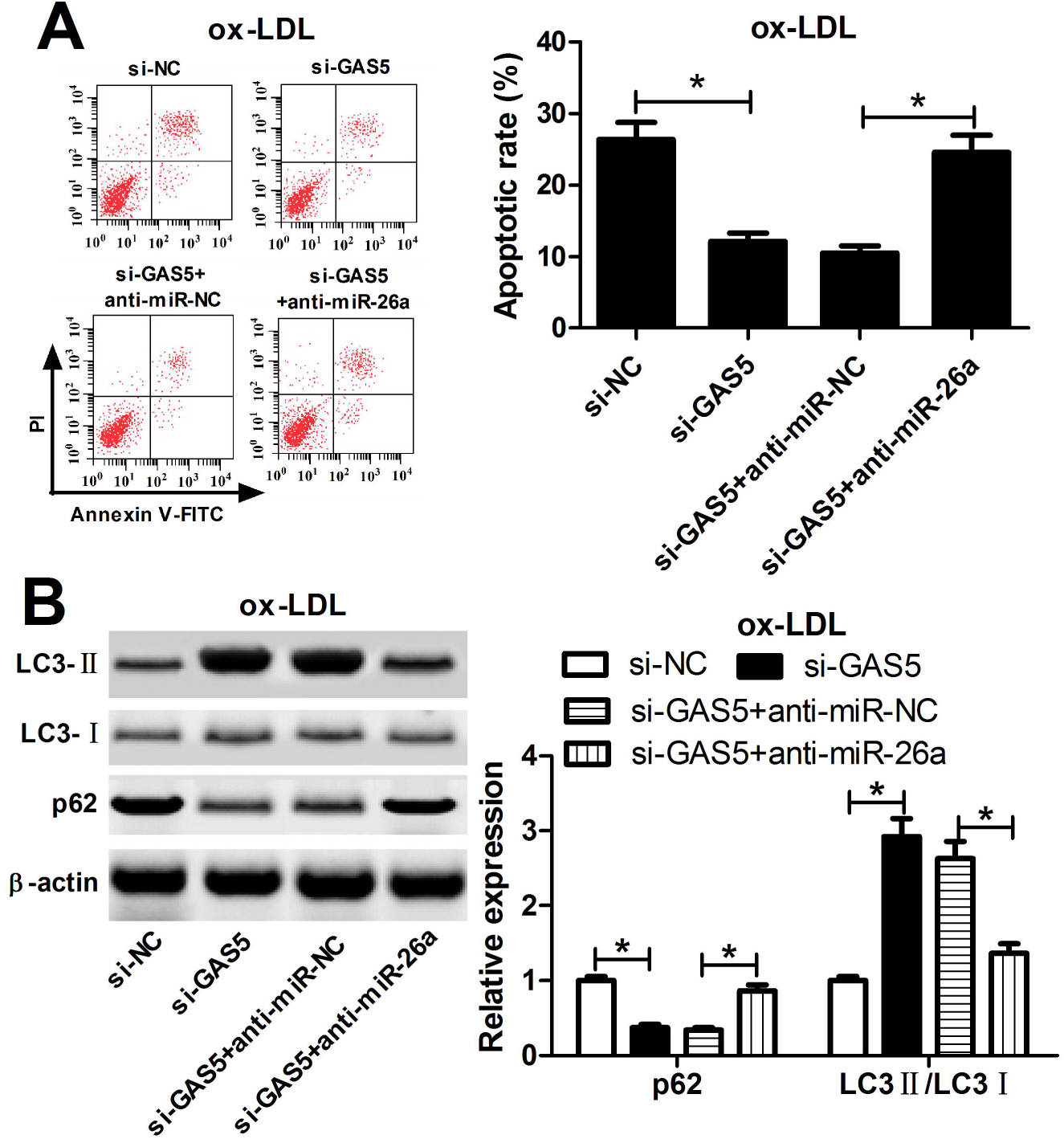
GAS5 knockdown restored ox-LDL-induced impaired autophagy flux by upregulating miR-26a in HAECs. HAECs were transfected with si-GAS5, si-NC, si-GAS5 + anti-miR-26a, si-GAS5 + anti-miR-NC, and then exposed to 1 mg/L ox-LDL for 24 h. (A) Flow cytometry analysis was performed to detect apoptosis of the treated HAECs. (B) Western blot was used to analyze the protein levels of LC3-II, LC3-I and p62 in the treated HEACs. \**P* < 0.05.

## 4 Discussion

In the present study, we demonstrated that GAS5 was upregulated and miR-26a was downregulated in the plasma samples of patients with atherosclerosis and ox-LDL-treated HAECs. Our study demonstrated that GAS5 directly interacted with miR-26a in ox-LDL-treated HAECs. Mechanistically, GAS5 knockdown restored ox-LDL-induced impaired autophagy flux by upregulating miR-26a in HAECs. Thus, GAS5 intervention may be a promising strategy to prevent atherosclerosis.

Dysregulation of autophagy has been documented to be closely associated with many diseases, including cardiovascular disease [21]. Autophagy is well-known to become dysfunctional in atherosclerosis, suggesting its protective role. It has been shown that the protective effect on autophagy might be beneficial to the therapy of atherosclerosis [22]. Recent studies showed that ox-LDL could induce vascular EC autophagy dysfunction and apoptosis in apoE^−/-^ mice [23]. Accordingly, we used ox-LDL to treat ECs and VSMCs to stimulate the pathological changes that occur during the early stage of atherosclerosis. During autophagy process, a cytosolic form of LC3 (LC3-I) is conjugated to form LC3-phosphatidylethanolamine conjugate (LC3-II) to promote autophagosome formation [24]. Thus, the ratio of LC3-II to LC3-I is considered as a marker for monitoring autophagy activity [25]. Additionally, p62, known as an autophagic substrate, is used as another widely marker of autophagy flux and can be selectively degraded by autophagy [26]. In our study, we found that ox-LDL stimulation induced apoptosis and impaired autophagy flux in HAECs, as demonstrated by the reduced ratio of LC3-II/LC3-I and increased expression of p62, which was consistent with the previous studies [27–29].

Recently, substantive studies have shown that ncRNAs including lncRNAs or miRNAs are identified as vital regulator of the physiological process of atherosclerosis [30]. For example, it was revealed that thymic stromal lymphopoietin (TSLP)-induced activation of lncRNA HOTAIR played a protective role in ox-LDL-induced EC injury by facilitating cell proliferation and migration and suppressing apoptosis in ECs [31]. In addition, miR-126 was reported to alleviate EC injury in atherosclerosis by restoring autophagic flux via inhibiting of PI3K/Akt/mTOR [32]. Recently, ample evidence has suggested that lncRNAs suppress the expressions and biological functions of miRNAs by acting as a ceRNA [33]. For example, it was demonstrated that silencing of H19 inhibited lipid accumulation and inflammation response in ox-LDL-treated Raw264.7 cells by upregulating miR-130b [34]. LncRNA H19 expression was found to be increased in atherosclerotic patient serum and ox-LDL-stimulated human aorta vascular smooth muscle cells (HA-VSMCs), and knockdown of H19 suppressed proliferation and induced apoptosis in ox-LDL-induced HA-VSMCs through modulating WNT/β-catenin signaling [35]. LncRNA MEG3 expression was reported to be downregulated in coronary artery disease (CAD) tissues than in normal arterial tissues, and ectopic expression of MEG3 increased EC proliferation and migration through inhibiting miR-21 expression [36].

In our study, we demonstrated that GAS5 was significantly upregulated and miR-26a was remarkably downregulated in the plasma samples of patients with atherosclerosis and ox-LDL-treated HAECs, which was in accordance with the previous studies [11, 12, 16, 17]. Given the inverse expression changes between GAS5 and miR-26a in atherosclerosis, we focused on the interaction between GAS5 and miR-26a. Luciferase reporter assay, RIP and RNA pull down manifested that GAS5 could directly interact with miR-26a by functioning as a ceRNA in ox-LDL-treated HAECs. Furthermore, rescue experiments demonstrated that we found that GAS5 knockdown alleviated ox-LDL-induced apoptosis and restored ox-LDL-induced impaired autophagy flux in HAECs by upregulating miR-26a.

## 5 Conclusions

In summary, our study provided the evidence that GAS5 knockdown restored ox-LDL-induced impaired autophagy flux by upregulating miR-26a in HAECs, revealing a novel regulatory mechanism for ox-LDL-induced impaired autophagy flux in ECs through ceRNA crosstalk. GAS5/miR-26a axis may extend our knowledge of the pathological mechanism of atherosclerosis and provided the potential therapeutic target for the treatment of atherosclerosis.

## Author Contributions

This work was designed and conceived by Weijie Liang. The experiment procedures and data analysis were carried out by Taibing Fan and Lin Liu. The manuscript was prepared by Weijie Liang and Lianzhong Zhang.

## Acknowledgements

Not applicable

ox-LDL: Oxidized low-density lipoprotein
EC: endothelial cell
GAS5: growth-arrest specific transcript 5
HAECs: human aortic endothelial cells
RIP: RNA immunoprecipitation
ncRNAs: noncoding RNAs
lncRNA: long noncoding RNA
miRNA: microRNA
HFD: high-fat diet
apoE: apolipoprotein E
ceRNA: competitive endogenous
RNA miR-NC: miRNA negative control
cDNA: complementary DNA
snRNA: small nuclear RNA
SDS-PAGE: sodium dodecyl sulfate-polyacrylamide gel
TBST: Tris-based saline with Tween 20
RISC: RNA-induced silencing complex
TSLP: thymic stromal lymphopoietin
HA-VSMCs: human aorta vascular smooth muscle cells
CAD: coronary artery disease

